# GplR1, an unusual TetR-like transcription factor in *Mycobacterium abscessus*, controls the production of cell wall glycopeptidolipids, colony morphology, and virulence

**DOI:** 10.1101/2025.05.07.652718

**Authors:** Scarlet S. Shell, Michal Bar-Oz, Junpei Xiao, Manitosh Pandey, Juan Bellardinelli, Mary Jackson, Stefan H. Oehlers, Daniel Barkan, Michal Meir

## Abstract

*Mycobacterium abscessus* is a major human pathogen, mostly infecting people with pre-existing lung conditions such as cystic fibrosis. The production of glycopeptidolipids (GPL) is a major determinant of virulence of this bacterium, with clinical isolates that lack GPL generally exhibiting more aggressive clinical behavior. The current paradigm is that GPL production is abolished *in vivo* via irreversible, spontaneous mutations taking place as part of in-host evolution. Little is known about the mechanisms or extent to which GPL production may be regulated. Here we describe an unusual TetR-like transcription factor of *M. abscessus*, MAB_1638, that appears to be a strong positive regulator of the entire GPL biosynthesis and export gene cluster through a combination of direct and indirect mechanisms. The inactivation of *mab_1638* abolished GPL production and thus led to stable rough colony morphology, as well as increased virulence in infection models, characteristic of rough, non-GPL-producers. Transcriptome analysis found the *mab_1638* mutant had 118 differentially expressed genes, including the GPL locus and a second, recently described GPL-like locus that produces a related glycosylated lipopeptide called GP8L. Chromatin Immunoprecipitation and sequencing revealed a consensus inverted-repeat DNA sequence motif characteristic of genes regulated by *mab_1638*. Together, *mab_1638* appears to encode a transcription factor required for production of GPL and therefore having a profound effect on virulence traits. We propose naming this gene **GPL regulator 1** (*gplR1*). This finding raises the important possibility that *M. abscessus* strains appearing smooth in laboratory growth conditions may nonetheless downregulate GPL-cluster genes in other conditions, including in-patient conditions, and thus acquire the phenotypic characteristics of rough strains.

**Importance:** *Mycobacterium abscessus* is an important human pathogen, causing disease that is difficult to treat. *M. abscessus* strains have been observed to have two distinct colony morphologies, smooth and rough, which substantially impact clinical presentation. Rough strains are associated with later-stage, more severe disease and are more virulent in animal models. Smooth morphology is conferred by a molecule called glycopeptidolipid in the outer cell envelope, and rough morphology is known to occur when mutations inactivate genes required for glycopeptidolipid biosynthesis. Little is known about the possibility that glycopeptidolipid production could be **regulated**. Here we have identified a transcription factor that is required for glycopeptidolipid biosynthesis, indicating that glycopeptidolipid production is indeed a regulated process, and raising the important possibility that strains exhibiting smooth morphology in the lab may down-regulated GPL production in the human host and thereby acquire the virulence properties of rough strains.

## Introduction

The genus *Mycobacterium* includes highly pathogenic species, such as *M. tuberculosis* and *M. abscessus*. These bacteria, as do others, use complex mechanisms to regulate their physiology and virulence. Bacterial genomes contain a variety of transcription factors that directly bind to regulatory sequences in the chromosome, most often in the promoters or upstream regulatory elements of the affected genes, resulting in higher or lower transcription. The identification of regulatory elements and their mechanisms of activity is important for understanding bacterial stress responses and pathogenesis.

An important virulence-associated mechanism in *M. abscessus* is the production of glycopeptidolipids (GPL), which are exported to the outer cell envelope. GPL production is a variable phenotype, evidenced by some clinical isolates but not others. Isolates producing GPL have a smooth colony morphology and, in some studies, have been reported to be more fit for intracellular growth and less pro-inflammatory than non-GPL producers (1, 2). Non-GPL producers have a rough morphology, cause more pronounced inflammation, and are associated with more severe and destructive disease (2-8).

It is widely accepted that GPL-negative isolates emerge *in-vivo* during chronic infection, and are associated with worse clinical outcomes (6, 9, 10). The majority of GPL-negative clinical isolates arise via inactivating mutations in the GPL-related genes – mostly the large non-ribosomal peptide synthase genes *mps1* and *mps2* (*mab_4099c* and *4098c*, respectively), and the transporter gene *mmpL4b* (*mab_4115c*) (10-12). However, some rough clinical isolates do not have mutations in these genes (6, 10, 12). Furthermore, strains that shift reversibly between smooth and intermediate morphologies have been observed (13), and our previous work demonstrated that a small regulatory RNA affects the expression of GPL biosynthesis and transport genes (14). Collectively, these observations suggest that in addition to all-or-nothing inactivating mutations, *M. abscessus* can regulate GPL production. We therefore opted to identify additional genes that are involved in the smooth-to-rough transition, possibly by regulating, rather than mutating, GPL-related synthetic pathways. Here we report the identification and characterization of a transcription factor required for high-level expression of GPL genes and thus GPL production and smooth morphology. This finding highlights the possibility that GPL production is not a constitutive phenotype for strains with WT GPL biosynthesis genes, but rather that GPL production is regulated in response to growth conditions.

## Results

### Transposon disruption of *mab_1638* leads to rough colony morphology due to loss of GPL production

We previously described the creation of a comprehensive transposon (Tn)-mutant library in *M. abscessus* (ATCC19977, smooth), based on a zeocin-selected Himar-1 transposon (15). We now turned to use this library to identify additional genes that lead to rough colony morphology when inactivated. We plated the Tn-library on 7H9/glycerol plates with no selection, picked several rough-appearing colonies, and re-plated them to confirm a stable rough phenotype. In those isolates with stable rough phenotypes, we identified the Tn-insertion sites as previously described (14). In one of these Tn-mutants, the identified insertion site was mapped to the TA at position 175 bp in the coding sequence of *mab_1638*. Given that the whole coding sequence was 648 bp, we expected this insertion to effectively inactivate the encoded protein. The insertion site was confirmed by PCR (Figure 1 a,b), followed by Sanger sequencing, and will subsequently be referred to as the *mab_1638*::tn strain. To confirm that the morphology phenotype was due to the inactivation of *mab_1638*, we complemented the *mab_1638*::tn strain with a single copy integrating vector containing an intact *mab_1638* coding sequence with the upstream 261 bp to include the native promoter. The complemented strain regained WT-like, smooth colony morphology (Figure 1c I-IV).

**Figure 1:**
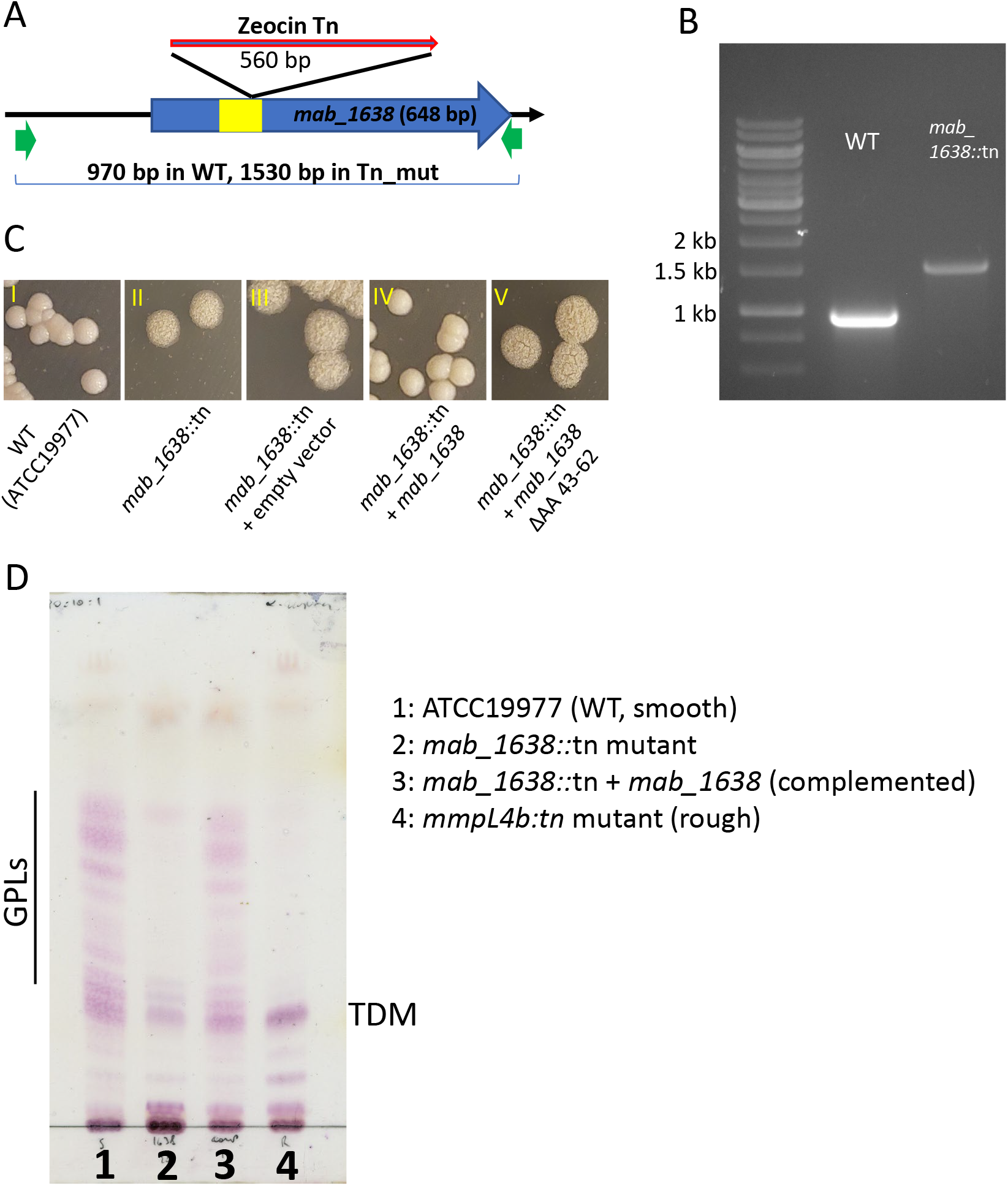
Disruption of mab_1638 causes rough morphology in *M. abscessus* that is due to lack of GPL in the cell envelope. **A**. Schematic representation of the transposon insertion location in ATCC19977. **B**. PCR confirmation of the insertion in mab_1638. **C**. Representative colonies of the indicated strains grown on 7H10 solid media. Images III-V show the *mab_1638::tn* insertion mutant transformed with the indicated plasmids. In the plasmids harboring *mab_1638*, the gene was expressed from its native promoter on an integrating plasmid. The plasmid in V expressed *mab_1638* with an in-frame deletion of the indicated amino acids. **D**. A TLC analysis of lipids extracted from the cell walls of the indicated strains. The predicted GPL bands are shown.

To examine whether the rough phenotype of the *mab_1638*::tn strain was due to lack of GPL in the cell envelope, we performed a TLC analysis of lipids extracted from the cell envelope of the mutant, as compared to the smooth, parental ATCC19977 strain and an *mmpL4b::tn* mutant with a classic rough morphology (Figure 1d). Whereas the smooth ATCC19977 strain had abundant GPL in the cell envelope, both the *mab_1638::tn* and *mmpL4b::tn* mutants did not have the characteristic bands. The complemented strain (lane 3) showed full reversion to the parental phenotype.

### MAB_1638 protein is a TetR-like transcription factor, with a DNA binding domain essential for its effect

*mab_1638* is designated in databases (such as Mycobrowser.epfl.ch) as a *tetR*-like transcription factor. By extrapolation from other TetR-like transcription factors, we identified the AA 43-62 of MAB_1638 as a helix-turn-helix structure that enables binding to DNA (16, 17). We therefore created a version of *mab_1638* with a deletion of AA 43-62, and introduced it into *mab_1638*::tn. In contrast to the intact version of the protein, MAB_1638^ΔAA43-62^ did not restore the WT-like, smooth colony morphology (Figure 1c, V).

### The inactivation of *mab_1638* in *M. abscessus* causes small increases in sensitivity to some antibiotics

TetR-family transcriptional regulators frequently repress expression of efflux pumps involved in antibiotic resistance, as has been shown previously in *M. abscessus* for MAB_2299c, MAB_2648c, and MAB_4384 (18-21). We therefore compared the drug sensitivity of the *mab_1638*::tn strain to its WT parent (ATCC19977). No significant differences in MIC were noted for meropenem, cefoxitin, trimethoprim/sulfamethoxazole, moxifloxacin and isoniazid (Table 1). However, the *mab_1638::*tn mutant had modestly decreased MIC values for vancomycin, teicoplanin, and linezolid as compared to the WT (Table 1 and Supplementary Fig. 1). The directionality of this effect is the opposite of what would be expected for a transcription factor that represses efflux pump expression. Additionally, these antibiotics have two distinct mechanisms of action; vancomycin and teicoplanin block peptidoglycan biosynthesis while linezolid blocks translation. A non-specific mechanism, such as changes in the myco-membrane composition and permeability (consistent with the colony morphology phenotype) may therefore be the cause for the subtle MIC shifts.

**Table 1:** The MICs (in µg/ml) of various antibiotics against *M. abscessus* strain ATCC19977 and its *mab_1638*::tn derivative.

| Drug | Drug Target | Method | ATCC19977 | *mab_1638::tn* | MIC fold change |
| --- | --- | --- | --- | --- | --- |
| Meropenem | Peptidoglycan (β-Lactam) | E-test | >32 | >32 | 1 |
| Cefoxitin | Peptidoglycan (β-Lactam) | E-test | 32 | 48 | 1.5 |
| Trimethoprim / sulfamethoxazole | Folate biosynthesis | E-test | >32 | >32 | 1 |
| Moxifloxacin | DNA gyrase | E-test | 1-2 | 1-2 | 1 |
| Vancomycin | Peptidoglycan (non β-Lactam) | E-test | 16 | 4 | 0.25 |
| teicoplanin | Peptidoglycan (non β-Lactam) | E-test | 12 | 3 | 0.25 |
| Linezolid | Ribosome | E-test | Poorly determined |  | 0.25 |
|  |  | Agar dilution | 50 | 12.5 |  |
| Isoniazid | Cell envelope (Mycolic acids synthesis) | Agar dilution | 2000 | 2000 | 1 |

### *M. abscessus mab_1638*::tn has a virulence phenotype resembling that of classic “rough” strains

To further characterize the phenotypic change conferred by *mab_1638* inactivation, we examined the mutant for virulence. We first used the adult zebrafish infection model of persistent granulomatous infection (22). Two groups of at least 8 adult zebrafish were infected with 10^7^ CFU of bacteria, and survival of the fish was followed for 10 days. Whereas there was no mortality in the group infected by WT bacteria, there was 50% mortality in the *mab_1638*::tn group by 3 days post infection and 100% mortality at day 9 (Figure 2a). An additional experiment was performed to assess CFU during infection. To allow longer survival, a lower inoculum of only 10^6^ CFU per fish was used (6 fish for WT infection, 8 fish for *mab_1638*::tn infection). However, even at this inoculum only half of the fish infected by the *mab_1638*::tn mutant survived until the sacrifice point on day 14. In those four surviving fish, no CFU difference was seen compared to the fish infected with the WT strain (Figure 2b).

**Figure 2:**
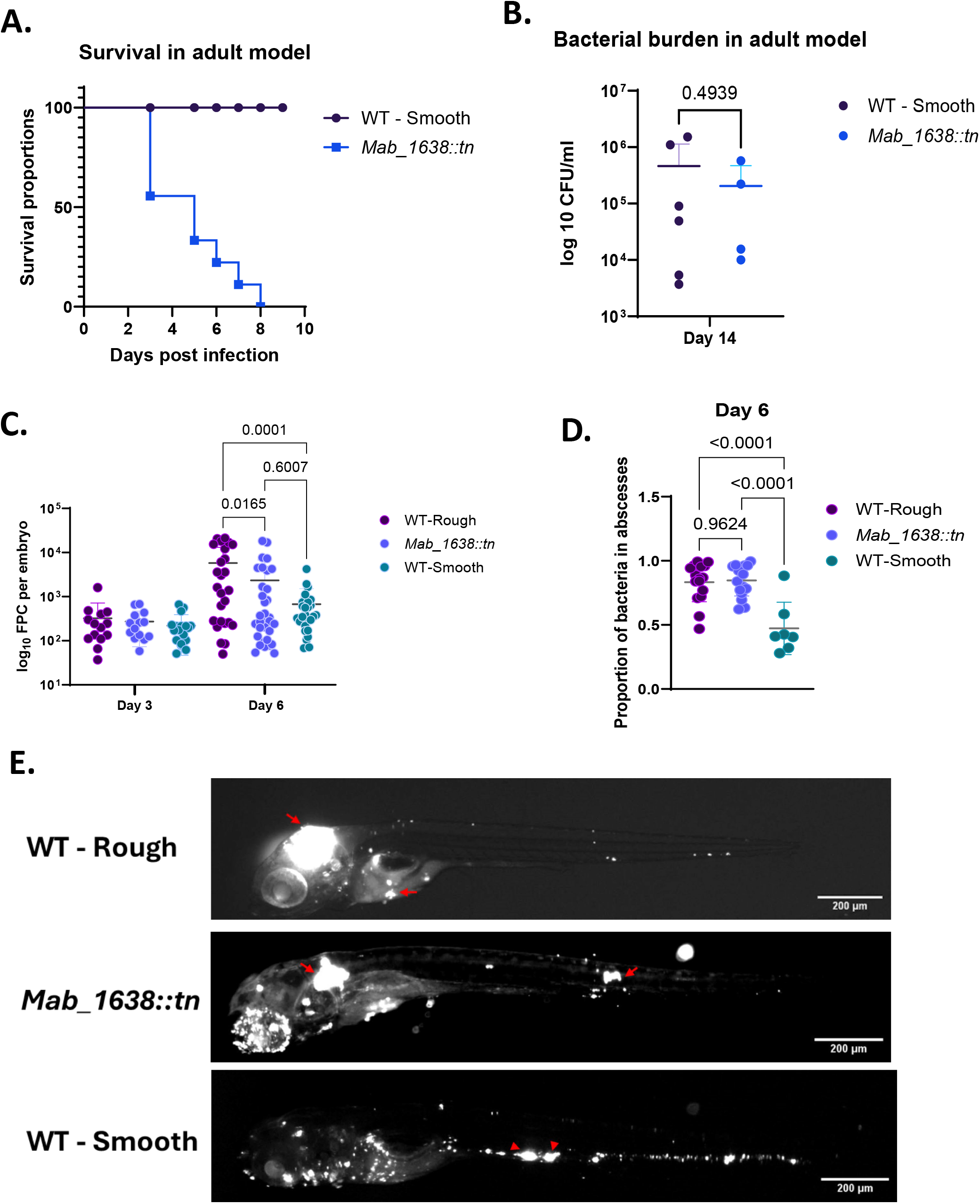
The *mab_1638:tn* mutant is hypervirulent in a zebrafish model and behaves similarly to a rough control strain. WT smooth is ATCC19977 and WT rough is ATC19977/CIP104536T-R. **A**: Adult zebrafish were infected by 107 cfu/fish, and survival was followed for 10 days. n=8 for WT, n=9 for *mab_1638:tn*. **B**: Bacterial burden was measured on day 14 post-infection by 106 cfu/fish and compared by unpaired t test. **C**: Bacterial burden, as represented by Fluorescent Pixel Count (PFC) was measured on day 3 and day 6 post infection in embryos infected by ∼250 cfu/embryo. **D**: The proportion of bacteria found in abscess-like structures was measured on day 6 post infection in embryos. Panels C and D, Ordinary one-way ANOVA with Tukey’s multiple comparisons test. **E**: Pictures of representative embryos from panel D. the red arrows point to clusters of bacteria (white fluorescence), referred to as “abscess-like”.

To determine if the *mab_1638*::tn strain had infection characteristics similar to those of a classic rough *M. abscessus* strain, we turned to live imaging of the zebrafish embryo-*M. abscessus* infection model, as our adult infection experiments yielded insufficient survivors for histological comparison. We performed an infection experiment with the *mab_1638*::tn mutant, smooth ATCC19977, and a rough variant of ATCC19977 called CIP104536^T^-R (2). Bacterial burden was measured on day 3 and day 6 post infection in embryos infected by ∼250 CFU/embryo. No difference in bacterial burden (as measured by fluorescence) was detected on day 3, but on day 6 there was a comparable increase in both the rough control and the *mab_1638*::tn mutant (Figure 2c). Consistent with previous reports, the rough control strain produced infections in which a greater proportion of bacteria were in large clusters (abscess-like structures) compared to the smooth control strain (Figure 2d-e). The behavior of the mutant was similar to that of the rough control strain, and distinct from that of the smooth parent strain (Figure 2d-e). Altogether, these results confirm that the *mab_1638*::tn mutant behaves like a GPL-null rough isolate, despite having no genetic defect in the GPL complex *per-se*.

### RNA sequencing reveals the transcriptional impact of *mab_1638*

To better understand the molecular mechanisms underlying the phenotypes of the *mab_1638*::tn strain, we first opted to characterize the transcriptome of the mutant as compared to WT bacteria by RNAseq. We grew both strains in independent triplicates, extracted RNA at similar growth-phase stages, and performed RNAseq as widely described. We found that 79 genes were significantly downregulated and 39 genes were upregulated (Figure 3a and Table S1; log_2_ fold change <-1 or >1 and adjusted *p* < 0.05). Strikingly, the downregulated genes included 18 genes in the GPL complex (*mab_4097c-4117c*), which are responsible for GPL production and export (Figure 3a). These downregulated genes included *mps1* and *mps2* (*mab_4099c* and *4098c*, respectively), as well as *mmpL4b*, all of which are implicated in the clinical transition of smooth to rough morphology (8, 10-12) and were downregulated by more than 8-fold. This downregulation may explain the strong rough phenotype of the *mab_1638*::tn strain, with no genetic disruption of the GPL locus itself.

**Figure 3.**
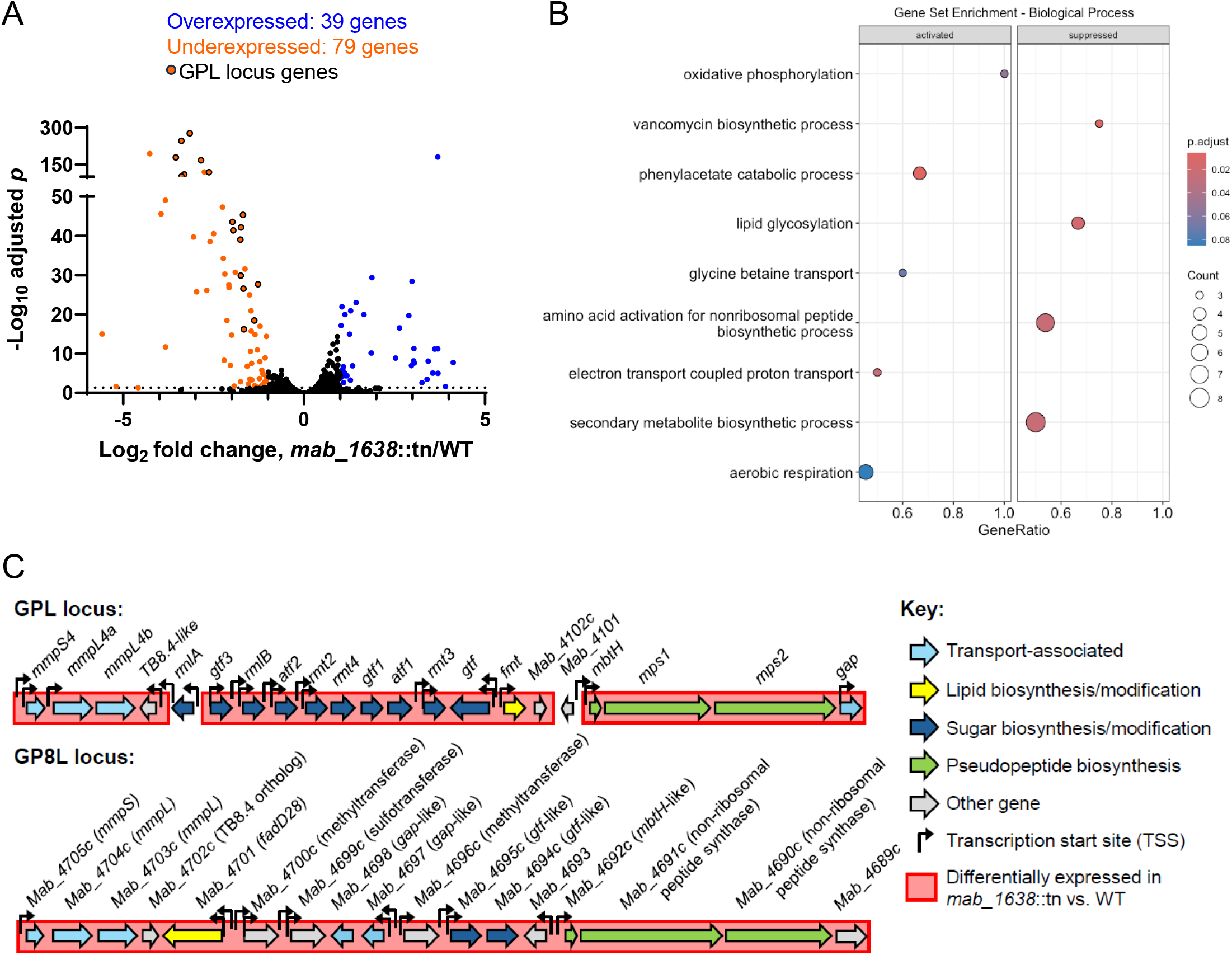
MAB_1638 negatively impacts expression of GPL biosynthesis and transport genes as well as the GP8L gene cluster. **A**. RNAseq revealed genes both up- and downregulated in the *mab_1638*::tn strain compared to the WT parental strain ATTC19977 (log_2_ fold change <-1 or >1, and adjusted *p* < 0.05). Genes known or expected to participate in GPL biosynthesis and transport are indicated. **B**. Gene set enrichment analysis was used to identify Gene Ontology Biology Process categories that were disproportionately affected by disruption of *mab_1638*. “Activated” genes had higher expression in the *mab_1638*::tn strain while “Suppressed” genes had lower expression. **C**. Diagrams of the GPL biosynthesis and transport gene cluster and a recently described GPL-like gene cluster termed the GP8L cluster that was also downregulated in the *mab_1638*::tn strain. Red shadowing indicates downregulated genes. Of the 38 total genes in both clusters, only 2 genes did not meet the criteria for downregulation, and are not shadowed. Genes are not shown to scale. Similarities between genes are indicated by naming and coloration. Bent arrows indicate reported TSSs.

Gene set enrichment analysis indicated that genes downregulated in the *mab_1638*::tn strain were enriched in several functional categories related to GPL biosynthesis (Fig. 3b; “lipid glycosylation,” “amino acid activation for nonribosomal peptide biosynthetic process,” and “secondary metabolite biosynthetic process”). Closer examination of the differentially expressed genes revealed the presence of a second locus spanning *mab_4689c-4705c* that included genes with similarities to the GPL locus genes, including predicted non-ribosomal peptide synthases, methyltransferases, sugar modification genes, and transport proteins (Fig. 3c). This GPL-related locus was recently reported to produce a glycosylated lipooctapeptide termed GL8P that is important for *M. abscessus* virulence in macrophage and mouse models (23). All of the genes in this GP8L locus were downregulated in the *mab_1638*::tn strain. The genes upregulated in the *mab_1638*::tn strain showed modest enrichment for those with roles in aerobic respiration, as well as enrichment for genes involved in phenylacetate metabolism. Finally, the differentially expressed genes included several predicted TFs, suggesting that some genes were directly controlled by MAB_1638 while others were affected indirectly.

### MAB_1638 recognizes an inverted repeat consensus sequence and appears to both positively and negatively regulate its targets

While the RNAseq results demonstrated the effect of *mab_1638* on the transcriptome, it was not clear which expression changes were due to direct binding of the MAB_1638 protein to its target genes and which were due to other transcription factors in the MAB_1638 regulon or downstream consequences of the physiological perturbations caused by changes in expression of MAB_1638’s targets. We therefore performed Chromatin Immunoprecipitation sequencing (ChIP-seq) to identify the sequences directly bound by MAB_1638. For the ChIP-seq, we cloned both N- and C-terminally HA-tagged versions of MAB_1638 separately in a multi-copy vector and inserted the resulting plasmids into the *mab_1638*:tn mutant. The N-terminally tagged version used a strong constitutive promoter while the C-terminally tagged version used the native promoter. We performed ChIP-seq with the two tagged strains and two negative control strains: WT (no HA tag), and a strain expressing an unrelated HA-tagged protein not expected to bind DNA. Genome regions with sequence coverage peaks were identified, and those present in both of the MAB_1638-tagged strains but absent from both negative controls were classified as putative MAB_1638 binding sites (Table S2). 324 of these putative binding sites were located within 500 nt upstream or downstream of a reported transcription start site (TSS) associated with a gene (24) (Table S3). We compared the frequencies of these putative binding sites among genes that were differentially expressed in the *mab_1638*:tn strain compared to the WT strain. Among genes with known TSSs, 30% of those that were downregulated and 53% of those that were upregulated had putative MAB_1638 binding sites, while only 15% of genes that were not differentially-expressed had putative MAB_1638 binding sites (Figure 4a). These statistically significant enrichments in putative binding sites among both up and downregulated genes suggest that MAB_1638 may act as both an activator and a repressor, depending on the target gene. While TetR-family TFs are often repressors, examples of activators in this family have been described (25, 26) as well as TetR-family proteins that serve as both activators and repressors (27-29).

**Figure 4.**
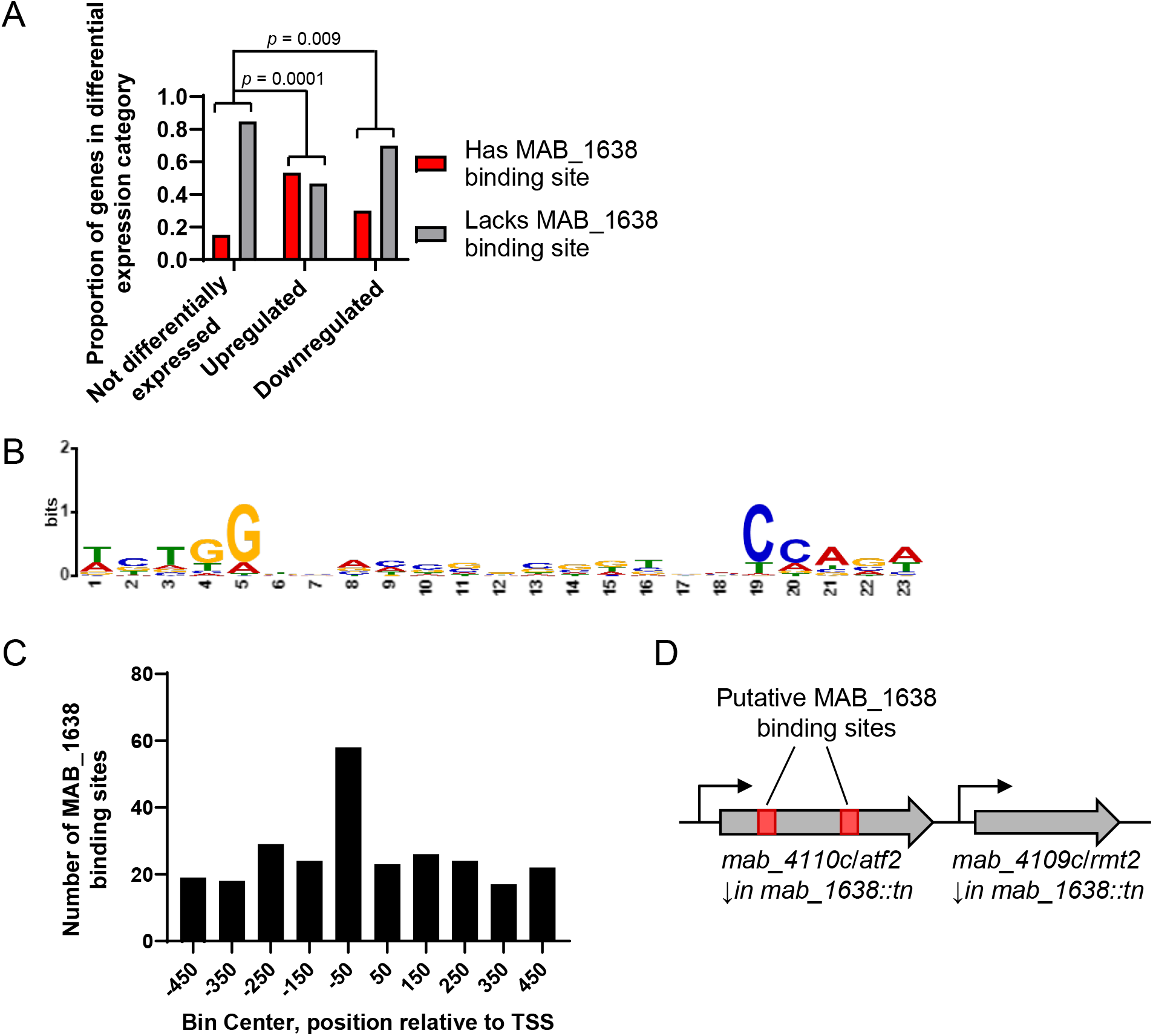
MAB_1638 binding sites are enriched in both up- and downregulated genes and feature an inverted-repeat consensus motif. **A**. ChIP-seq was used to identify genomic sites bound by MAB_1638. Genes with annotated TSSs were then stratified based on the presence or absence of MAB_1638 binding sites within 500 nt of their TSSs and how their expression was affected by disruption of *mab_1638*. Genes that were up- or downregulated in the *mab_1638*::tn strain were significantly more likely to have MAB_1638 binding sites in their putative regulatory regions than genes that were not differentially expressed (Fisher’s exact test). **B**. MEME analysis revealed an inverted repeat motif present in ∼80% of the MAB_1638 binding sites in putative regulatory regions. **C**. MAB_1638 binding sites motifs in putative regulatory regions (motif within 500 nt up- or downstream of an annotated TSS) were disproportionately located in the regions ∼100 bp upstream of TSSs. **D**. Two inverted-repeat MAB_1638 binding site motifs were identified within the GPL biosynthesis locus. Their approximate locations are indicated.

Motif analysis with MEME and FIMO (30) revealed an inverted repeat motif in 266 (82%) of the 324 putative MAB_1638 binding sites associated with TSSs (Figure 4b). The motif was 23 nt in length, consisting of 11 nt inverted repeat arms separated by a single nt. The outer 5 nt of each repeat had a more conserved sequence (5’-TCTGG-3’) than the inner portion. This motif was disproportionately frequent in the 100 nt upstream of TSSs (Figure 4c), consistent with expectations for a typical bacterial TF. Although most of the GPL biosynthesis and transport genes were downregulated in the *mab_1638*::tn strain, there were only two MAB_1638 binding sites detected within the GPL gene cluster, both in the coding sequence of *mab_4110c* (*atf2*) and upstream of the TSS of *mab_4109c* (*rmt2*) (Figure 4d). Like most GPL genes, both *atf2* and *rmt2* were downregulated in the *mab_1638*::tn strain. Collectively, these results indicate that MAB_1638 is a bona fide transcription factor that binds an inverted repeat motif to regulate expression of various genes including some involved in GPL biosynthesis. MAB_1638-binding motifs were detected for genes that were both up- and downregulated in the *mab_1638*::tn strain, consistent with the idea that MAB_1638 acts as both an activator and as a repressor.

### Defining the MAB_1638/GplR1 regulon

While 118 genes were differentially expressed in the *mab_1638*::tn strain according to the cutoffs described above (Table S1), many of these genes are likely to be indirectly affected and therefore not technically part of the MAB_1638 regulon. Those differentially expressed genes for which ChIP-seq revealed MAB_1638 binding (23 genes, defined by having a binding peak within 500 nt of an annotated TSS) are likely to be direct targets. However, this gene list is limited to those genes with annotated TSSs (65/118 differentially expressed genes). Genes lacking TSSs may plausibly be encoded in polycistronic operons together with upstream genes that have annotated TSSs. We therefore considered differentially expressed genes lacking TSSs to be likely members of the MAB_1638 regulon if they were encoded immediately downstream of differentially expressed genes with TSSs and MAB_1638 binding sites. This added eight additional genes, expanding the putative MAB_1638 regulon to include 31 genes in total (Table 3).

Given the substantial impact of *mab_1638* on GPL gene expression and the likelihood that additional transcription factors regulating GPL gene expression will be discovered, we propose naming it GPL regulator 1 (*gplR1*).

## Discussion

Bacteria regulate their genetic and metabolic networks, and subsequently virulence, by a complex network of DNA-binding proteins – transcription factors, that in turn are generally responsive to various environmental cues. In *M. abscessus*, a major virulence factor (or arguably, an anti-virulence factor) are the glycolpeptidolipids (GPL), which when present in the outer cell envelope give the bacterial colony its smooth dome shape. Lack of GPL in the cell envelope confers a rough colony morphology, and leads to more destructive inflammatory responses, extracellular growth, resistance and persistence to some antibiotics – collectively causing a more virulent phenotype with a considerably worse prognosis (6-10, 31, 32). It is thought that patients are usually infected by smooth strains, and via *in-vivo* evolution acquire rough isolates that drive the pathological clinical picture. Some rough variants arise via inactivating mutations in GPL producing or transporting genes (the cluster *mab_4096* through *mab_4117*), but many rough clinical isolates have what appear to be intact GPL-cluster genotype (6, 10, 13), suggesting there are regulatory pathways that can up- or down-regulate GPL production and transport, perhaps in response to environmental cues. We recently identified and published one such regulatory pathway involving the sRNA B11 (14), where inactivation of B11 was sufficient to cause a rough phenotype. We therefore postulated that additional factors may regulate, rather than simply inactivate, GPL production/export, and here we have described one of these factors.

We used a transposon-mutant library to identify a mutant with a transposon insertion in the previously unstudied gene *mab_1638*, which exhibited a stable rough morphology on agar plates. This mutant recapitulated the hyper-virulent phenotype associated with “classic” rough isolates when tested in the zebrafish model (adult and embryo). The protein encoded by *mab_1638* is predicted to have the classic structure of a TetR-like transcription factor – a large family of TFs present in most bacteria and involved in multiple physiologic and pathogenesis processes. Indeed, when we complemented the Tn-mutant with a version of the gene coding for a protein unable to bind DNA, smooth colony morphology was not restored, supporting the hypothesis that DNA binding is required for the protein to exert its effect.

The combination of transcriptomic and ChIP-seq analyses provided the landscape of the *mab_1638* regulon. 31 genes were differentially expressed and had putative GplR1/MAB_1638 binding sites within their likely regulatory sequences, suggesting they are direct targets. This putative regulon likely includes some false positives, as GplR1/MAB_1638 appeared to bind the regulatory regions of ∼15% of non-differentially expressed genes (Fig. 4a). There are also likely additional regulon members that we did not detect because their TSSs have not yet been annotated. None of the *gplR1/mab_1638* regulon genes are located near *gplR1/mab_1638* on the *M. abscessus* chromosome. This makes GplR1/MAB_1638 a Type III TetR family transcriptional regulator according to a scheme published by Ahn and colleagues (33). Unlike more conventional TetR family regulators which regulate a small number of adjacently transcribed genes, Type III regulators control disparately located sets of genes.

The closest homolog of GplR1/MAB_1638 in *M. tuberculosis* is Rv0275c, which was previously described as mediating isoniazid resistance in *M. bovis* (34). However, *M. abscessus* encodes another predicted TetR regulator, MAB_2594, that has more similarity and identity to Rv0275c than GplR1/MAB_1638 does. It is therefore unsurprising that disruption of *gplR1/mab_1638* did not affect isoniazid resistance (Supplementary Fig. 1).

The putative GplR1/MAB_1638 regulon defined here includes 8 genes that appear to be negatively regulated by GplR1/MAB_1638 and 23 genes that appear to be positively regulated (Table 2). This contrasts with previously described TetR family transcription factors in *M. abscessus*, MAB_2299c, MAB_2648c, MAB_4384, which all negatively regulate drug efflux pumps (18-21). Additional mycobacterial TetR family regulators have been described that negatively regulate cholesterol metabolism genes (35, 36), CRISPR genes (37), oxidative stress response genes (38, 39), among others. There is a precedent in *M. smegmatis* for TetR family regulators that directly activate expression of their targets; *msmeg_6564* was reported to activate expression of DNA repair genes (40), and *msmeg_4022* was reported to activate expression of drug transporters (41). To our knowledge, the mechanisms of positive regulation in those cases are not known. To understand how GplR1/MAB_1638 carries out both positive and negative regulation, it will be necessary to experimentally test the binding sites of GplR1/MAB_1638 in its putative target promoters, the impact of *gplR1/mab_1638* on the activity of these promoters, and the TSSs involved.

**Table 2.** The putative regulon of MAB_1638 direct targets. Genes differentially expressed in the *mab_1638*::tn strain were included if they had MAB_1638 binding peaks or were likely to be co-transcribed with genes with MAB_1638 binding peaks.

| Gene | Annotated TSS<sup>a</sup> | Plausibly in operon with MAB_1638 target<sup>b</sup> | MAB_1638 binding peak | Log<sub>2</sub> fold change *mab_1638::tn*/ WT<sup>c</sup> | P adjusted<sup>c</sup> | Uniprot annotation |
| --- | --- | --- | --- | --- | --- | --- |
| MAB_0852 | yes | NA | yes | -2.3 | 4.5 x 10<sup>-48</sup> | Possible conserved polyketide synthase associated protein PapA2 |
| MAB_0853 | no | yes | no | -2.2 | 5.7 x 10<sup>-35</sup> | Probable esterase/lipase LipP |
| MAB_0854 | no | yes | no | -2.2 | 5.1 x 10<sup>-31</sup> | Fatty acyl-AMP ligase FadD28 and polyketide synthase |
| MAB_0855 | no | yes | no | -1.9 | 1.8 x 10<sup>-31</sup> | Membrane protein, MmpL family |
| MAB_0884c | no | yes | no | -2.5 | 2.6 x 10<sup>-41</sup> | Mycobacterium 19 kDa lipoprotein antigen |
| MAB_0885c | yes | NA | yes | -4.3 | 1.7 x 10<sup>-194</sup> | Hypothetical lipoprotein lpqH |
| MAB_0939 | yes | NA | yes | 1.7 | 4.9 x 10<sup>-75</sup> | Probable polyketide synthase |
| MAB_1013 | yes | NA | yes | -2.8 | 1.2 x 10<sup>-73</sup> | Ribonucleases G and E |
| MAB_1042c | yes | NA | yes | 3.9 | 0.024 | Cytochrome c oxidase subunit 1 |
| MAB_1312 | yes | NA | yes | -1.2 | 1.0 x 10<sup>-17</sup> | Beta-lactamase-like |
| MAB_1313 | no | yes | no | -1.2 | 1.0 x 10<sup>-08</sup> | Probable transcriptional regulator, AraC family |
| MAB_1565c | yes | NA | yes | 1.1 | 1.1 x 10<sup>-20</sup> | Alpha-amylase family protein |
| MAB_1674c | yes | NA | yes | -2.1 | 3.5 x 10<sup>-19</sup> | Amidase AmiA2 |
| MAB_1827c | yes | NA | yes | -1.2 | 0.00024 | Bacteriophage protein |
| MAB_2464 | yes | NA | yes | -1.5 | 0.0053 | Chitin-binding type-2 domain-containing protein |
| MAB_2734c | yes | NA | yes | 1.3 | 1.1 x 10<sup>-15</sup> | Lipoprotein |
| MAB_2735c | yes | NA | yes | 1.9 | 7.2 x 10<sup>-11</sup> | Helix-turn-helix domain-containing protein |
| MAB_2741 | yes | NA | yes | 1.4 | 1.0 x 10<sup>-23</sup> | DUF5666 domain-containing protein |
| MAB_3448 | yes | NA | yes | -1.4 | 0.00050 | Acetyltransferase |
| MAB_4109c | yes | NA | yes | -1.4 | 3.5 x 10<sup>-19</sup> | Methyltransferase |
| MAB_4110c | yes | NA | yes | -1.7 | 1.3 x 10<sup>-30</sup> | Probable acetyltransferase AtfA |
| MAB_4438 | yes | NA | yes | 1.7 | 1.1 x 10<sup>-20</sup> | Serine aminopeptidase S33 domain-containing protein |
| MAB_4469c | yes | NA | yes | -1.0 | 4.3 x 10<sup>-15</sup> | Sulfatase family protein |
| MAB_4689c | no | yes | no | -3.1 | 1.8 x 10<sup>-40</sup> | Uncharacterized protein |
| MAB_4690c | no | yes | no | -2.9 | 3.5 x 10<sup>-67</sup> | Probable non-ribosomal peptide synthetase PstA |
| MAB_4691c | no | yes | no | -3.4 | 6.1 x 10<sup>-94</sup> | Probable non-ribosomal peptide synthetase PstA |
| MAB_4692c | yes | NA | yes | -5.6 | 1.0 x 10<sup>-15</sup> | MbtH-like protein |
| MAB_4693 | yes | NA | yes | -2.1 | 1.5 x 10<sup>-27</sup> | Probable cytochrome P450 |
| MAB_4775c | yes | NA | yes | -1.5 | 1.2 x 10<sup>-21</sup> | L,D-transpeptidase |
| MAB_4778c | yes | NA | yes | -1.3 | 1.2 x 10<sup>-11</sup> | Uncharacterized protein |
| MAB_4891c | yes | NA | yes | 1.9 | 4.1 x 10<sup>-30</sup> | HTH lysR-type domain-containing protein |

<sup>a</sup>As reported in (24).
<sup>b</sup>Based on absence of a reported TSS and examination of coverage in our RNAseq data.
<sup>c</sup>DESeq2

One of the largest groups of genes transcriptionally affected by *gplR1/mab_1638* disruption was a set of 18 genes from the GPL locus, half of which were downregulated at least 8-fold, thus explaining the rough colony morphotype in the absence of genetic mutations in the GPL locus itself. However, only two GplR1/MAB_1638 binding sites were identified in the GPL locus, both within the *atf2* coding sequence. One of these binding sites is located ∼484 nt upstream of the annotated TSS of the downstream gene, *rmt2*, and could plausibly contribute to the regulation of its expression. The other binding site is located >1.2 kb upstream of the annotated *rmt2* TSS and therefore seems unlikely to regulate its expression. Further work will be needed to determine if GplR1/MAB_1638 binding at these sites impacts of expression of *rmt2* or other genes in the GLP locus. It appears likely that at least some of the impact of *gplR1/mab_1638* on GLP gene expression is indirect, given the absence of GplR1/MAB_1638 binding sites elsewhere in the locus. Notably, several predicted transcription factors were differentially expressed in the *mab_1638::*tn strain, and two of these had GplR1/MAB_1638 binding sites revealed in our ChIP-seq data. Additional transcription factors under the control of GplR1/MAB_1638 may therefore be direct regulators of GPL locus genes.

Interestingly, a second large group of genes with similarity to the GPL locus genes were also downregulated. These were recently reported to encode proteins that produce GP8L, a glycosylated lipooctapeptide (23). GP8L is clearly not sufficient to independently confer smooth morphology in laboratory conditions, since various works have shown that single mutations in key canonical GPL locus genes cause rough morphology. Like canonical GPL, the role of GP8L in virulence appears to be complex. A strain in which the GP8L locus was disrupted in a rough background had reduced uptake by macrophage-like THP-1 cells, and possibly also a growth defect following uptake (23). In zebrafish embryo infections, a rough strain with GP8L disruption produced a bacterial burden lower than the rough parental strain and similar to a smooth strain (23). Finally, the rough GP8L disruption strain caused less mortality in mice than its rough parent (23). The impact of GP8L on strains that also produce GPL (smooth strains) is unknown, as is the significance of the apparent co-regulation of GPL and GP8L production by GplR1/MAB_1638. These questions should be investigated further.

Mounting evidence – both from examining clinical isolates (6, 12, 13) and from targeted studies like this one and our previous study on the B11 sRNA (14) - point to the conclusion that “S to R” transitions are not limited to random mutagenesis events selected by *in-vivo* evolutionary pressures, but that the bacteria has active mechanisms to regulate GPL production and thus associated phenotypes. This is consistent with previous work in *M. smegmatis* showing that outer cell envelope composition is subject to transcriptional regulation (42). Thus far we have identified the *gplR1/mab_1638* gene and the B11 sRNA, both of which appear to maintain GPL production under certain conditions (including those in the lab), but may, in response to some as-yet unidentified cue, downregulate GPL production and export to produce the more virulent phenotype we associate with the “rough morphology”. When these bacteria are isolated from patients or from infection models and re-grown on agar plates, this phenotype would logically reverse to “smooth colony” in response to the unknown signals in the lab growth environment that lead to expression of positive regulators of GPL such as *gplR1/mab_1638* and B11. This raises the question whether in clinical practice, many of the so called “smooth isolates” may actually behave in a “rough” isolate manner in the patient, but that only a tip of the iceberg of these isolates is identified as “rough” when re grown in the clinical microbiology lab. This means research should focus on the regulatory elements governing the GPL formation, and the metabolic cues that may signal the bacteria to downregulate it. Interfering with these signals may provide a yet-unexplored therapeutic pathway.

## Methods

### Strains and growth conditions

All strains were derived from *M. abscessus* ATCC19977 WT (smooth). Bacteria were grown in Middlebrook 7H9 supplemented with ADS (final concentrations

### Isolation, identification, and subsequent complementation of a *mab_1638::*tn mutant

The creation of a zeocin-selected transposon-mutant library in *M. abscessus* was previously described (15). The identification of the insertion site was performed the same way as the library sequencing, only on a single colony. The primers used for corroboration of the insertion site (Fig 1a) were 1638-F: 5’ – CAGAGCCATGCCTGGATCCAG – 3’ and 1638-R: 5’ – AGCGTGCTCAGTGCTCGACGG -3’. The *mab_1638*::tn strain was also referred to as mDB292. For complementation, the same primers were used to PCR-amplify a 970 bp fragment from *M. abscessus* gDNA (containing the gene and the 261 upstream bases), and cloned into the SwaI site of the attB-integrating vector pDB213, to create pDB426. To create the ΔAA43-62 mutant, PCR was used to create two separate fragments, one ending at AA 42 and the other starting at AA 63, and then fusing the two fragments by the above described 1638-F and 1638-R primers to get a 910 bp fragment with the bases coding for AA 43-62 deleted. This fragment was again cloned into the SwaI site of pDB213, to create pDB448.

### Total lipid extraction and TLC analysis

*M. abscessus* ATCC19977 WT (smooth), *mab_1638*::tn, *mab_1638*::tn + *mab_1638*, and *mmpL4b*::tn (rough) were grown in 7H9 ADS Tw 0.05% until OD = 0.4. Cells were collected, washed with PBS and lipids extracted with CHCl_3_:CH_3_OH (1:2 vol/vol) once, followed by CHCl_3_:CH_3_OH (2:1 vol/vol) twice. After Folch wash, lipids were weighed, and the same amount of lipids were loaded onto a silica gel 60 F254 coated aluminum sheet. Samples were developed in CHCl_3_:CH_3_OH:H_2_O (90:10:1) and revealed with α-naphthol.

### Antibiotic sensitivity and MIC determination

For E-tests, equal amounts of the tested bacteria were plated on 7H9/glycerol plates in top (soft) agar to ensure equal spread of bacteria, and an E-test strip (BD BBL™ Sensi-Disc™, Becton Dickinson and Company) was placed. Plates were incubated at 37°C, and results were read after 96-120 hours. Isoniazid sensitivity was performed using agar dilution: 7H9/glycerol/ADS plates were supplemented by the determined concentrations of isoniazid, and the tested bacteria were inoculated using a 10 µl loop from a 0.02 OD_600_ culture.

### Zebrafish infection models

Fluorescent labeling was done by mWasabi and mTomato expressing plasmids (a kind gift from Kevin Takaki and Lalita Ramakrishnan, Cambridge University). Zebrafish infection experiments were approved by A*STAR IACUC in protocol 221694. *M. abscessus* WT and *mab_1638*::tn fluorescently labeled strains were grown in 7H9 broth supplemented with 0.5% glycerol, 10% OADC and 0.05% Tween 80 in the presence kanamycin 250µg/ml. Single cell suspension stocks for bacterial infection were prepared according to a previously reported protocol (22). Briefly, 0.8 OD_600_ cells were centrifuged and washed with 1x PBS and resuspended in 7H9 broth. Cells were declumped by passaging through a 27G needle three times and stored at -80°C in aliquots.

Adult zebrafish from AB strain were infected intraperitoneally with fluorescently labelled *M. abscessus* WT and *mab_1638*::tn strains using a 31G needle. Infected zebrafish were housed in a recirculating aquarium system (28°C, 750 µS conductivity, pH 7.4) with a 14/10 hour light/dark cycle and observed as part of their daily feeding routine (22).

CFU recovery was performed on euthanized adult zebrafish by homogenization in a bead beater and plating on LB agar for 4-5 days growth at 37°C.

Zebrafish embryos were infected by microinjection (43). Embryos were anesthetized, mounted in 3% methyl cellulose, and imaged at 6 dpi on a Nikon SMZ25 equipped with a Nikon Digital Camera DS-10. Image analysis was performed with FIJI (ImageJ) to enumerate bacterial fluorescent pixel count and abscesses were defined as having an area of 200 px corresponding to approximately 500 mm^2^ (1.60 mm /px) (3).

Strains used as controls were the ATCC19977 (Smooth) and the ATC19977/CIP104536^T^-R (rough).

### RNA sequencing and analysis

*M. abscessus* WT and *mab_1638*::tn were grown in triplicates (each starting from a different single colony) to an OD_600_ of 0.5 in 7H9 supplemented with 0.5% glycerol and 0.05% Tween 80. Bacteria were collected and RNA was extracted as previously described (14). Briefly, frozen cultures were thawed on ice and centrifuged at 4,000 rpm for 5 min at 4°C. The pellets were resuspended in 1 ml Trizol (Life Technologies) and placed in tubes containing Lysing Matrix B (MP Bio). Cells were lysed by bead-beating twice for 40 s at 9 m/sec in a FastPrep 5G instrument (MP Bio). RNA was purified using Direct Zol RNA miniprep kit (Zymo) according to manufacturer’s instructions. rRNA depletion, library preparation and RNA sequencing were performed in the Technion Genomics Core Facility (Haifa, Israel). Data are available on GEO, accession number GSE287636.

Reads were aligned to the NC_010397 reference genome using the Burrows-Wheeler Aligner (44). The FeatureCounts tool (45) was used to assign mapped reads to genomic features, and DESeq2 (46) was used to assess changes in gene expression. Genes with adjusted p-values < 0.05 and log^2^ fold changes > 1 or < -1 were considered to be differentially expressed.

The R package clusterProfiler 4.0 (47) was used to perform pathway enrichment analysis using KEGG annotation data. clusterProfiler was also used to conduct gene set enrichment analysis (GSEA) using Gene Ontology (GO) annotations from QuickGO (https://www.ebi.ac.uk/QuickGO/annotations) (48).

### Chromatin immune-Precipitation (ChIP) sequencing and analysis

For N-terminal tagging, an HA-tag sequence was added to the 5’ end of the *mab_1638* coding sequence, and this fragment was cloned under the strong constitutive MOP promoter. For C-terminal tagging, the HA-sequence was simply added to the 3’ end of *mab_1638* driven by its native promoter. Both constructs were cloned into the episomal multi-copy vector pDB32 to create pDB458 and pDB459, respectively. These were introduced into the *mab_1638*::tn mutant to create strains mDB353 (MOP promoter + N-terminal tag) and mDB352 (native promoter + C-terminal tag). Two negative control strains were assayed in parallel: WT ATCC19977, and a strain expressing an HA-tagged cyclopropane synthase (MmaA2) from *M. tuberculosis*, which is expected not to bind DNA. Immunoprecipitation was performed based on the method described in (49). Bacteria were cultured overnight in 50 mL of 7H9 medium supplemented with 0.5% glycerol and 0.05% Tween 80 to an OD_600_ of ∼0.4-0.6, and equilibrated to room temperature. Formaldehyde (1% final concentration) was added, followed by a 20-minute shaking-incubation at room temperature. Cross-linking was quenched by adding glycine (final concentration of 0.25 M) for 5 minutes at RT. Fixed cultures were collected by centrifugation at 4500 x g for 10 minutes. Pellets were washed twice with cold PBS and resuspended in 600 µL of ChIP lysis buffer (20 mM HEPES-KOH pH=7.9, 50 mM KCl, 0.5 mM DTT, and 10% glycerol) supplemented with a protease inhibitor cocktail (Roche cOmplete #11836153001). The resuspended cells were then transferred to tubes containing 500-700 mg of 0.1 mM glass beads (Sigma-Aldrich G8772). Bead beating was performed through three cycles, each consisting of 30 seconds of beating followed by a 5-minute pause on ice, at 4260 RPM. Samples were centrifuged at 100 x g for 2 minutes at 4°C, and supernatants were collected. DNA shearing was achieved using a Covaris M220 instrument with the following parameters: 10% duty cycle, 10 minutes, and 200 cycles per burst. Insoluble materials were removed by centrifugation at 11,000 x g for 10 minutes at 4°C, and the supernatant was collected. Efficiency of shearing was confirmed by electrophoresis on a 2% agarose gel. Anti-HA antibodies (3 µg per sample, Abcam, Ab#9110) were added to 25 µg of sheared DNA from each sample, followed by a 1-hour incubation at 4°C with rotation. Subsequently, 25 µL of pre-washed magnetic beads (NEB-S1425S) were added to the samples, and incubation was continued overnight with rotation at 4°C. The next day, the samples were subjected to magnetic separation to remove liquids. The beads were washed five times with 900 µL of IPP50 buffer (10 mM Tris-HCl buffer pH=8, 150 mM NaCl, and 0.1% NP-40), followed by two washes with TE buffer (50 mM Tris-HCl pH=8, 10 mM EDTA). Beads-antibody-DNA complexes were collected by magnetic separation and then resuspended in 70 µL of elution buffer (50 mM Tris-HCl pH=8, 1% SDS). This suspension was incubated at 65°C for 15 minutes. The elution step was repeated with additional 70 µL, and the eluents were combined. RNA was removed from the eluted DNA using RNase A (NEB T3018-2, 20 mg/mL) at 37°C for 1 hour. Subsequently, reverse crosslinking was carried out by adding 3 µL of proteinase K (20 mg/mL, MAN-740506) and incubating for 2 hours at 37°C, followed by an overnight incubation at 65°C. The next morning, DNA was extracted using phenol:chloroform. Finally, the extracted DNA was resuspended in 500 µL of 10 mM Tris-HCl pH=8.

Library preparation, sequencing, quality control, and peak analyses were conducted by the Technion Genomics Center, Haifa, Israel. 100 bp single-end reads were aligned to *Mycobacteroides abscessus* reference genome (https://ftp.ncbi.nlm.nih.gov/genomes/all/GCA/000/069/185/GCA_000069185.1_ASM6918v1/GCA_000069185.1_ASM6918v1_genomic.fna.gz) and annotation file (https://ftp.ncbi.nlm.nih.gov/genomes/all/GCA/000/069/185/GCA_000069185.1_ASM6918v1/GCA_000069185.1_ASM6918v1_genomic.gff.gz) using Bowtie2 v2.2.6 with default parameters, filtering reads for unique alignment using samtools v1.3. Peaks detection was done using MACS2 v2.1.1. Data are available on GEO, accession number GSE287715.

Peaks were filtered to identify those that were present in samples from both strains expressing tagged MAB_1638 and absent in the two negative control strain samples. Each peak had start and end coordinates as well as a peak summit coordinate. The start and end coordinates were used to define peaks as being present in both MAB_1638-tagged strains, with > 90% overlap required. Peaks that passed this filter were considered to be putative MAB_1638 binding sites (Table S2). Putative MAB_1638 binding sites were classified as associated with a gene if the peak summit was within 500 nt upstream or downstream of a TSS annotated as gene-associated in (24) (Table S3). For gene-associated putative MAB_1638 binding sites, we extracted 25 nucleotides upstream and 25 nucleotides downstream of the peak summit position. These sequences were input into MEME to discover de novo motifs (30) using the *M. abscessus* genome composition as the background. The discovered motif was then used as input for FIMO (50) to identify other potential motifs within the whole peak region. This step helps locate additional instances of the discovered motif, since the peak summit may not always represent the exact location of the MAB_1638 binding site.

## Acknowledgements and funding

D.B is supported by the Israeli Science Foundation IPMP-grant (1349/20) and an individual grant (608/22). S.H.O and M.P were supported by the Singapore NMRC, under its OF-IRG program, Award OFIRG22jul-0081 to S.H.O. S.S.S and J.X. were supported by NIH/NIAID grant 1P01AI143575-01A1 to S.S.S. M.M. is supported by the Israeli Science Foundation 1730/22, and the CF Foundation (https://www.cff.org/) award number 005596I223.

## Data availability

RNAseq data and ChIPseq data are available on NCBI GEO, accession numbers GSE287636 and GSE287715, respectively.

## Supplemental Materials

<u>Figure S1</u>: *Mycobacterium abscessus* WT (ATCC19977) and the *mab_1638::tn* mutant (mDB292) were tested by E-test for the MIC to vancomycin (A), teicoplanin (B), and by agar dilution to linezolid (C) and isoniazid (D). In (C,D), bacteria were streaked on 7H9/glycerol agar plates with the designated concentrations of linezolid (1.5-50 µg/ml) or isoniazid (500-2000 µg/ml).

<u>Figure S2:</u> For the ChiP seq, the *mab_1638::tn* mutant (rough) was transfected with tagged version of the native 1638 gene, with an HA-tag attached either at the N or the C terminus of the protein. To demonstrate the tagged protein is functional, single colonies were examined for morphology. As seen, both tagged versions of 1638 revert the colony morphology back to smooth, confirming proper function.

<u>Table S1</u>. Transcriptomics comparison of *M. abscessus* ATCC19977 and *gplR1*/*mab_1638*::tn.

<u>Table S2</u>. Putative GplR1/MAB_1638 binding sites identified by ChIP-seq.

<u>Table S3</u>. Putative GplR1/MAB_1638 binding sites associated with genes and their sequence motifs.

