## Supplemental Figures for "GplR1, an unusual TetR-like transcription factor in *Mycobacterium abscessus*, controls the production of cell wall glycopeptidolipids, colony morphology, and virulence"

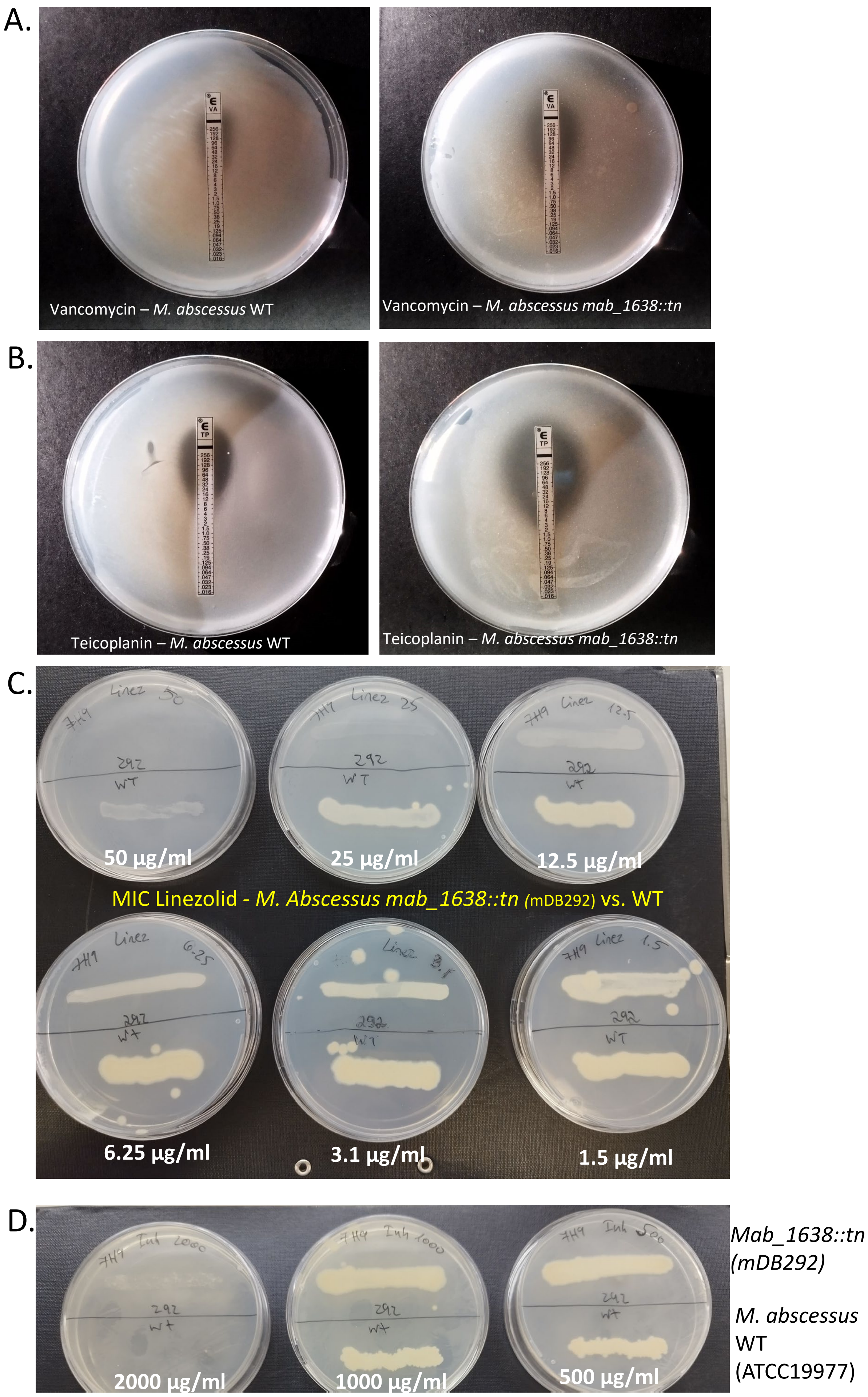


<u>**Supplementary Figure 1**</u>: *Mycobacterium abscessus* WT (ATCC19977) and the *mab_1638::tn* mutant (mDB292) were tested by E-test for the MIC to vancomycin (A), teicoplanin (B), and by agar dilution to linezolid (C) and isoniazid (D). In (C,D), bacteria were streaked on 7H9/glycerol agar plates with the designated concentrations of linezolid (1.5-50 µg/ml) or isoniazid (500-2000 µg/ml.

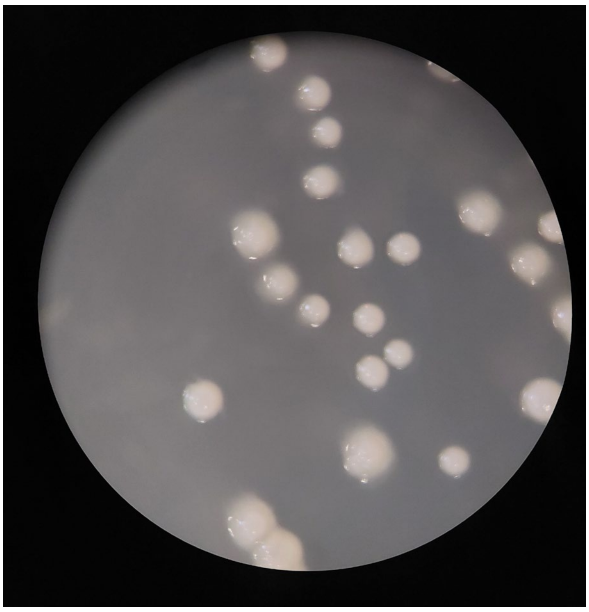

mDB352 (*mab_1638*::tn mutant + 1638-HA tag). The tag is located at the C-terminus of the protein.

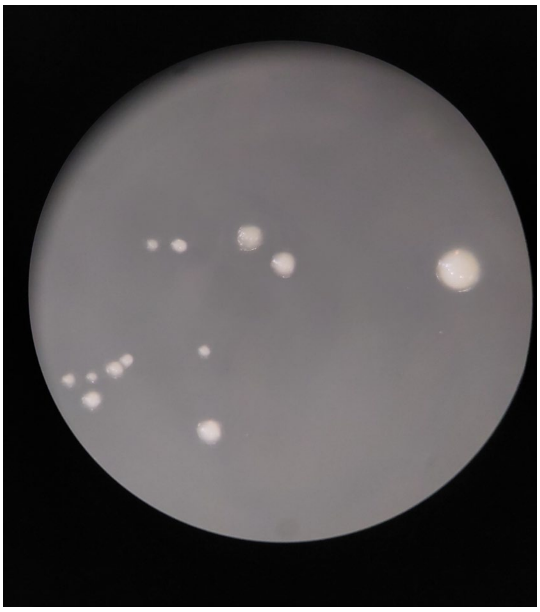

mDB353 (*mab_1638*::tn mutant + $HA_{tag}$-1638). The tag is located at the N-terminus of the protein.

**Supplemental Figure 2**: For the ChiP seq, the *mab_1638*::tn mutant (rough) was transfected with tagged version of the native *mab_1638* gene, with an HA-tag fused either to the N or the C terminus of the protein. To demonstrate the tagged protein is functional, single colonies were examined for morphology. As seen, both tagged versions of MAB_1638 revert the colony morphology back to smooth, confirming proper function.
